# Differential gene expression by RNA-Seq in Sigma-2 Receptor/TMEM97 knockout cells reveals its role in complement activation and SARS-CoV-2 viral uptake

**DOI:** 10.1101/2021.03.14.435180

**Authors:** Aladdin Riad, Yann Aubert, Chenbo Zeng, Thomas J. A. Graham, E. James Petersson, Brian C. Capell, Robert H. Mach

## Abstract

Our lab has recently shown that the Sigma-2 Receptor/Transmembrane Protein 97 (sigma-2R/TMEM97) interacts with the low-density lipoprotein receptor (LDLR) and facilitates the enhanced uptake of various ligands including lipoproteins and intrinsically disordered proteins. TMEM97 has been recently been shown to interact with severe acute respiratory syndrome coronavirus 2 (SARS-CoV-2) viral proteins, highlighting its potential involvement with viral entry into the cell. We hypothesized that sigma-2R/TMEM97 may play a role in facilitating viral uptake, and with the regulation of inflammatory and thrombotic pathways that are involved with viral infection. In this study, we identified the top differentially expressed genes upon the knockout of sigma-2R/TMEM97, and analyzed the genes involved with the inflammatory and thrombotic cascades, effects that are observed in patients infected with SARS-CoV-2. We found that the ablation of sigma-2R/TMEM97 resulted in an increase in Complement Component 4 Binding Protein (C4BP) proteins, at both the translational and transcriptional levels. We also showed that sigma-2R/TMEM97 interacts with the cellular receptor for SARS-CoV-2, the human angiotensin-converting enzyme 2 (ACE2) receptor, forming a protein complex, and that disruption of this complex results in the inhibition of viral uptake. The results of this study suggest that sigma-2R/TMEM97 may be a novel therapeutic target to inhibit SARS-CoV-2 viral uptake, as well as to decrease inflammatory and thrombotic effects through the modulation of the complement cascade.

## INTRODUCTION

Despite being implicated in the pharmacology of several well-known agents that have assisted in elucidating the fundamental biology of the sigma-2 receptor/TMEM97, the receptor’s biological function remains to be fully elucidated. Sigma receptors, which consist of two subtypes termed sigma-1 and sigma-2 receptors, are a distinct family of proteins and are present in the periphery and central compartment ^1-3^. Following initial radioligand binding studies of the sigma-2 receptor, Xu, *et al*. observed with photoaffinity labeling that the sigma-2 receptor resided within a protein complex containing progesterone membrane binding component 1 (PGRMC1). ^4^. Work by Kruze and colleagues, TMEM97 was identified as the gene that encodes the sigma-2 receptor; this work led to the renaming of the protein as the sigma-2R/TMEM97 ^5^. Subsequent studies from our group identified the presence of a protein complex between sigma-2R/TMEM97 and PGRMC1 ^6^, studies which set the foundation for the pharmacological characterization of this protein complex. For example, our lab showed that sigma-2 receptor/TMEM97 and PGRMC1 interact with the LDLR, and the three form a trimeric protein complex that is essential for the efficient uptake of lipoproteins ^6^. Using CRISPR/cas9 technology, we demonstrated that the loss or pharmacological inhibition of sigma-2R/TMEM97 resulted in the decreased uptake of lipoproteins such as low-density lipoprotein (LDL) ^7^. Sigma-2R/TMEM97 also has been shown to be involved with cholesterol homeostasis and metabolism through its interaction with the cholesterol transport-regulating protein Niemann-Pick C1 (NPC1) and through the sterol response element-binding protein (SREBP) pathway ^6,8,9^.

Recently, our lab has shown that the sigma-2 receptor/TMEM97 is involved in processes including the uptake of apolipoprotein E (apoE) and intrinsically disordered proteins via its interaction with the LDL-receptor-related protein (LRP) ^6,7^. These studies were conducted using the sigma-2 receptor KO cell line described above or with sigma-2R/TMEM97 inhibitors in cultured neurons^6^. More recently, the sigma-2R/TMEM97 has been identified as a binding protein for severe acute respiratory syndrome coronavirus 2 (SARS-CoV-2) viral protein Orf9c and has been identified as having the potential to be a therapeutic target for the inhibition of viral infectivity ^10^. SARS-CoV-2 viral entry has been shown to be mediated via the recognition of the viral spike protein with the host receptor angiotensin-converting enzyme 2 (ACE2) along with the assistance of serine protease Transmembrane Serine Protease 2 (TMPRSS2) ^11^. SARS-CoV-2 ORF9c Is a membrane-associated protein that suppresses antiviral responses in cells ^12^. Complications with coronavirus disease 2019 (COVID-19) comprise a wide variety of symptoms such as increased inflammation including the overactivation of the complement system, and these complications can be related to stroke, thrombotic events, and immune system overactivation in the afflicted patients ^13-15^.

The goal of the current study was to use our sigma-2R/TMEM97 k/o cells to further explore the cell biology of this protein. In the first set of experiments, we conducted RNA-seq analysis to would further shed light on the biological pathways interacting with the sigma-2 receptor. Our results have identified a previously unrecognized association between the receptor coagulation and complement pathways. A second goal of this study was to determine if sigma-2R/TMEM97 is involved in SARS-CoV-2 viral entry into cells. In these studies, we identified an interaction between sigma2/TMEM97 and ACE2 receptor and characterized the receptor’s role in viral uptake into human cells. Taken together, our studies have revealed a novel role of the sigma-2R/TMEM97 in SARS-CoV-2 viral entry into sells as well as in modulating the coagulation cascade and the complement system that are frequently observed in COVID-19. Our results highlight sigma-2R/TMEM97’s potential as a unique receptor that may prove to be an important therapeutic target for preventing or treating COVID-19 or other infections by the coronaviruses^16-18^.

## RESULTS

### RNA-Seq analysis on HeLa sigma-2R/TMEM97 KO cells

HeLa sigma-2R/TMEM97 KO cells were plated and RNA-Seq was performed to analyze differentially expressed genes compared to scramble/cas9 control cells (Fig 1). The top 25 upregulated and downregulated genes showed several genes that play a role in molecular pathways involving cancer development, which was no surprise as sigma-2R/TMEM97 has been shown to be a proliferation marker and has been studied extensively in the context of cancer ^19-23^.

**Figure 1.**
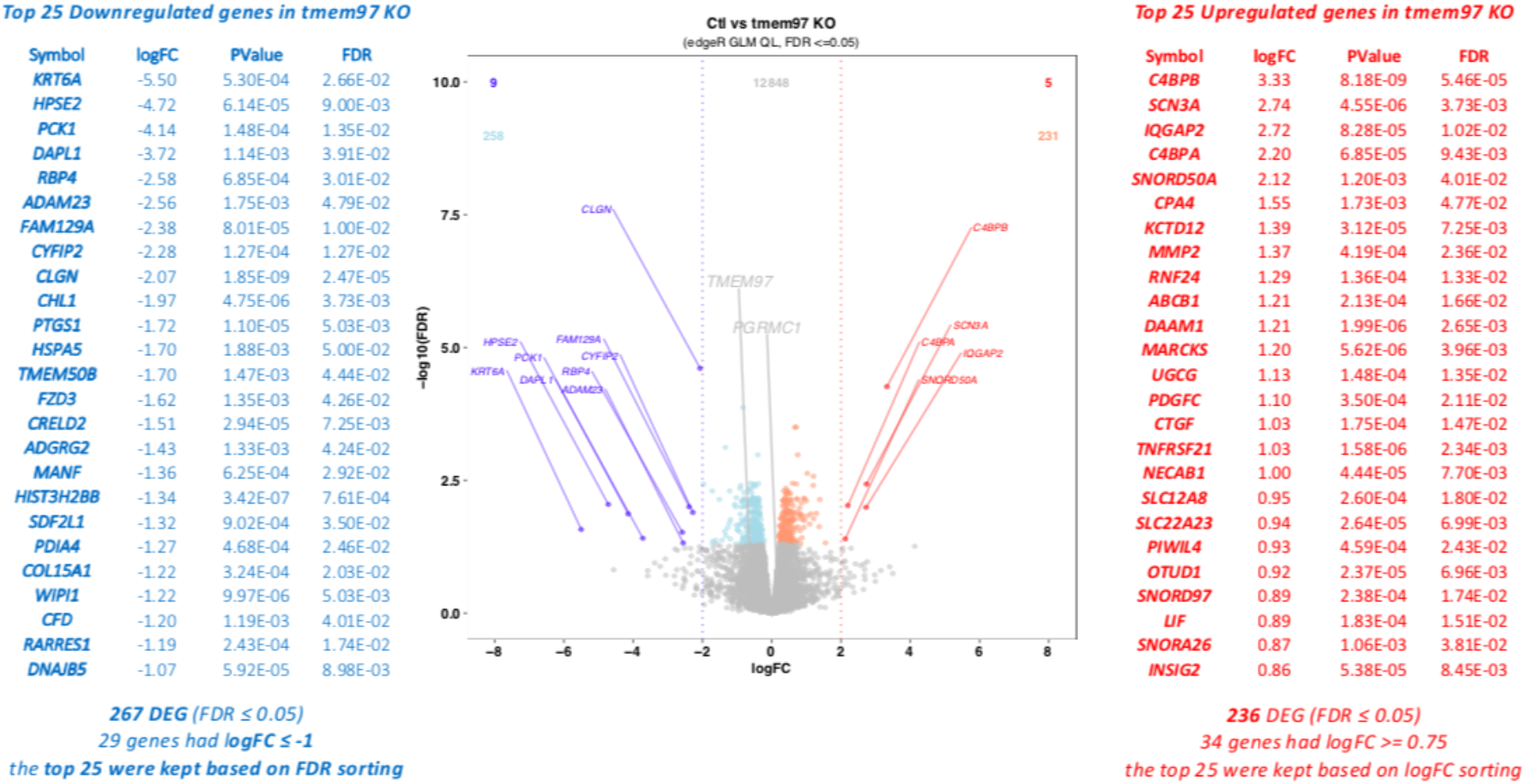
Top Differentially Expressed Genes in HeLa TMEM97 KO Cells. RNA-Seq was performed on scramble/Cas9 and TMEM97 KO HeLa cells and the top 25 differentially expressed genes were identified (FDR≤0.05; logFC≤-1 for downregulated genes; logFC≤0.85 for upregulated genes)

Interestingly, Complement Component 4 Binding Protein (C4BP) Beta (C4BPB) and Complement Component 4 Binding Protein Alpha (C4BPA) were amongst the top upregulated genes upon loss of sigma-2R/TMEM97. C4BP is involved in the inflammatory pathway through the inactivation of the complement cascade and is involved in coagulation and thrombosis pathways through its interaction with Protein S ^24-26^. C4BP has also been shown to play a role in inflammation triggered by other viruses such as Influenza A Virus (IAV) ^27^. For these reasons, we were interested in understanding the role of sigma-2R/TMEM97 in these pathways as inflammation and coagulation have been shown to play significant roles in COVID-19 ^28-33^.

### Quantification of secreted C4BP in HeLa model system using ELISA

Our next objective was to verify that the genetic expression levels observed via RNA-Seq corresponded with an increase in C4BP at the protein level. As C4BP is a secreted protein, HeLa scramble/cas9 and sigma-2R/TMEM97 KO cells were plated for 72 hours, and the C4BP protein level was quantified in the media (Fig 2A) via ELISA. The C4BP levels secreted by sigma-2R/TMEM97 KO cells were significantly increased compared to those of control cells (Fig 2A), verifying that the upregulation of C4BP at the transcriptional level corresponded with an increase in translation. To assess if this increase in secreted C4BP could be induced using highly selective sigma-2R/TMEM97 ligands, scramble/cas9 cells were plated in the presence of increasing concentrations of RHM-4. IL-6 was used as a positive control as it has been shown to increase the secretion of C4BP ^34-36^. RHM-4, which is selective to sigma-2R/TMEM97, increased the secretion of C4BP in a had a dose-dependent manner (Fig 2B), indicating the specific pharmacological modulation of sigma-2R/TMEM97 increases secreted C4BP protein similar to what was observed through modulation at the transcriptional level. Taken together, these results indicated that sigma-2R/TMEM97 is a potential therapeutic target for the inhibition of complement system activation, and may play a role in the modulation of the coagulation cascade through the upregulation of C4BP.

**Figure 2.**
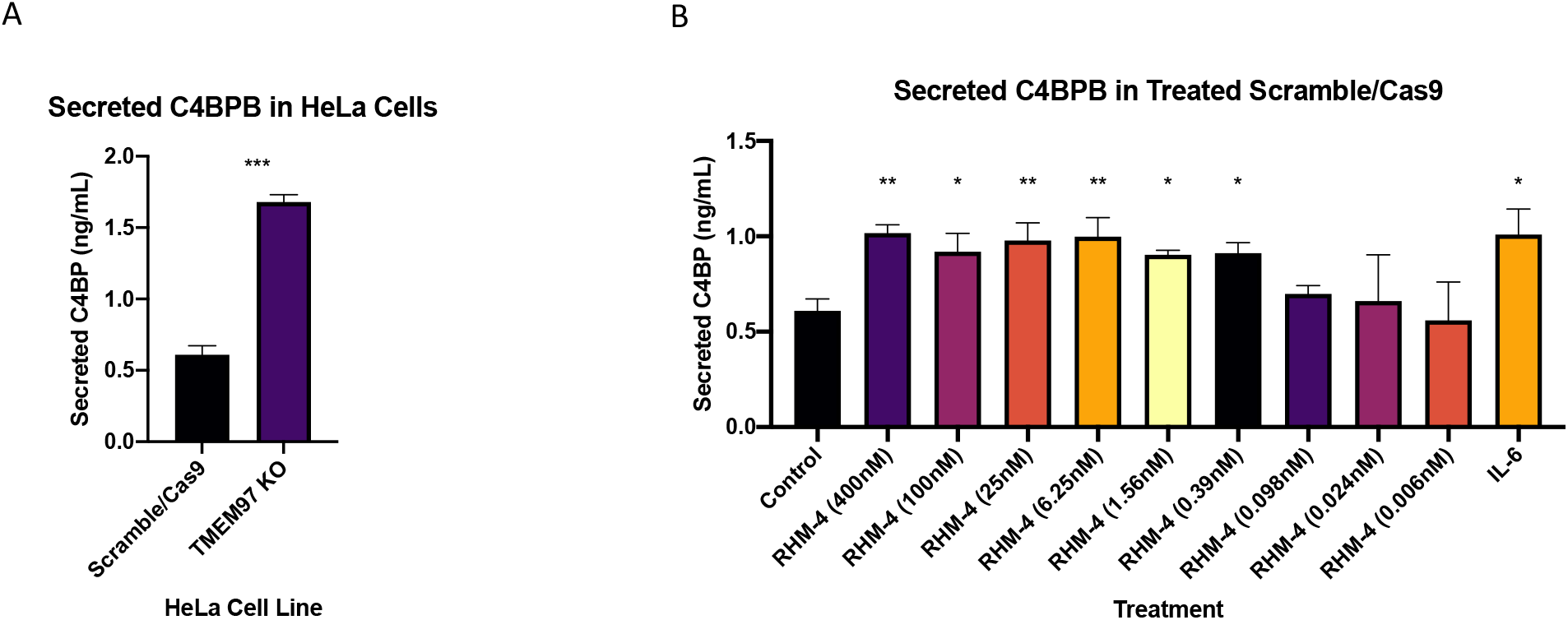
C4BPB Secretion is increased in cells with ablated TMEM97. C4BPB was measured using an ELISA 72 hours post treatment. (A) TMEM97 KO HeLa cells showed significantly increased C4BPB secretion compared to control Scramble/Cas9 cells. (B) Treatment of scramble/cas9 HeLa cells with RHM-4 showed an increase in C4BPB in a dose-dependent manner, treatment with 5ng/mL IL-6 was used as a positive control. (n=3, * p ≤ 0.05; ** p≤0.01; *** p≤0.001)

### Loss of sigma-2R/TMEM97 results in inhibition of SARS-CoV-2 pseudovirus uptake

Our next goal was to assess role of sigma-2R/TMEM97 in the uptake of a pseudovirus that expresses the SARS-CoV-2 spike protein. HeLa scramble/cas9 control cells and sigma-2R/TMEM97 KO cells were plated, transduced with human ACE2, and then treated with the SARS-CoV-2 pseudovirus (Fig 3 A and B). Upon transduction with ACE2, sigma-2R/TMEM97 KO cells showed a significant reduction in the uptake of SARS-CoV-2 pseudovirus (Fig 3A). As expected, without human ACE2, expression of sigma-2R/TMEM97 had no significant effect upon pseudovirus uptake (Fig 3B). These sigma-2R/TMEM97 data indicated that both sigma-2R/TMEM97 and human ACE2 were necessary for efficient uptake of the virus, the absence of either protein resulted in ineffective pseudovirus uptake. The loss of sigma-2R/TMEM97 reduces pseudovirus uptake to the basal level seen in cells lacking human ACE2. Furthermore, RHM-4 and haloperidol treatment of scramble/cas9 control cells that have been transduced with human ACE2 showed an inhibitory effect on viral uptake (Fig 3C). As RHM-4 is specific to sigma-2R/TMEM97, this study was important in showing that specifically targeting sigma-2R/TMEM97 results in the inhibition of viral uptake, as haloperidol is a non-selective sigma receptor ligand, whereas RHM-4 is specific for the sigma-2R/TMEM97, indicating that the specific inhibition of sigma-2R/TMEM97 attenuates pseudovirus uptake. This inhibition was not observed in cells that were not ACE2 competent (Fig 3D), indicating that it’s inhibitory role is intimately tied with the ACE2 receptor’s function of viral recognition. Collectively, these findings showed the role of sigma-2R/TMEM97 in enhancing viral uptake in an ACE2-dependent manner.

**Figure 3.**
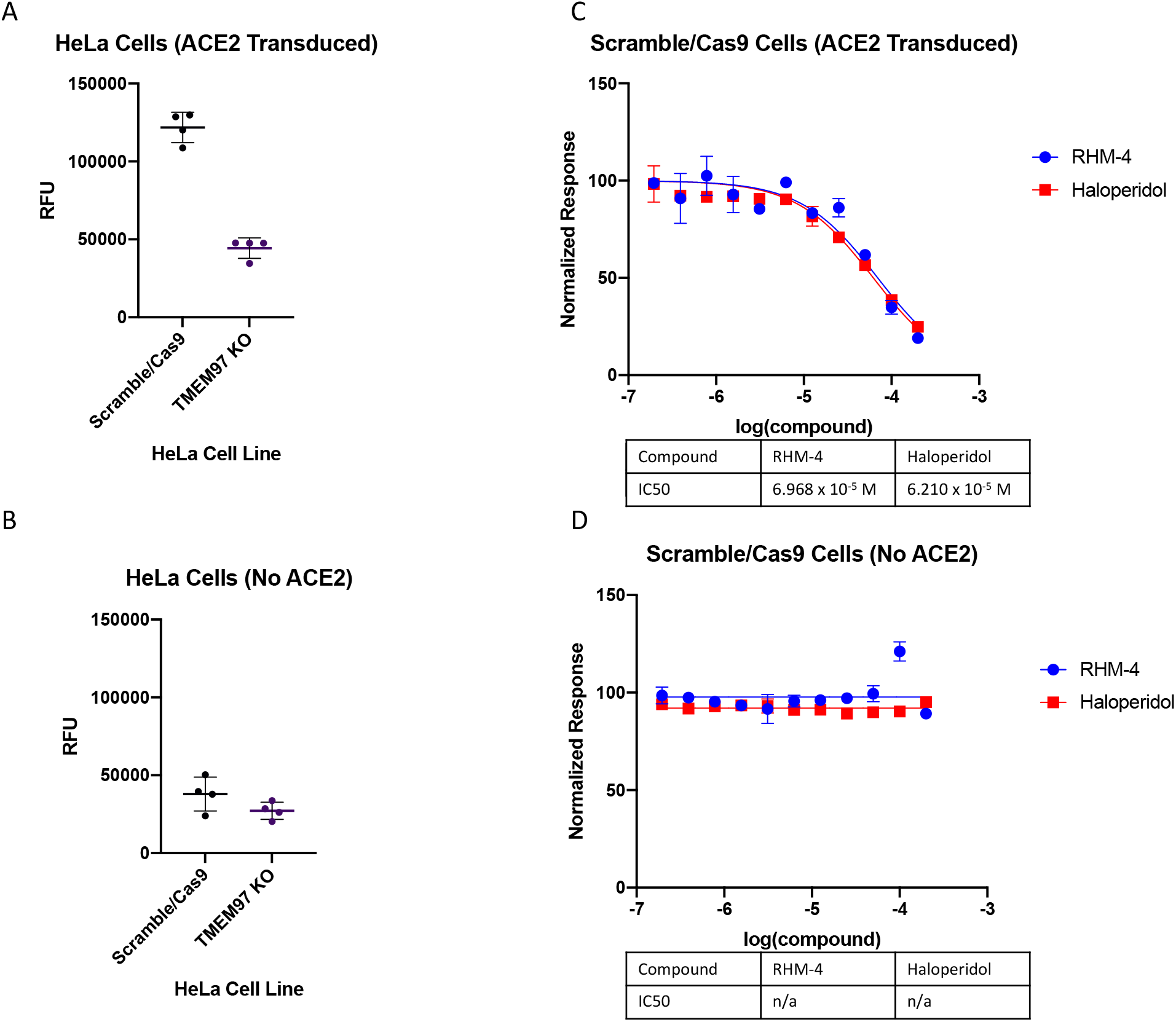
TMEM97 inhibits SARS-CoV-2 pseudovirus uptake in ACE2 competent cells. (A) TMEM97 KO HeLa cells transduced with human ACE2 showed reduced uptake of fluorescent SARS-CoV-2 pseudovirus particles over 24 hours compared to (B) cells that were not transduced with human ACE2. (C) Scramble/Cas9 HeLa cells treated with RHM-4 or Haloperidol showed a dose dependent reduction in SARS-CoV-2 pseudovirus. (D) Cells that were not transduced with human ACE2 did not display an effect. (n=3, * p ≤ 0.05; ** p≤0.01; *** p≤0.001)

### Sigma-2R/TMEM97 forms a complex with the human ACE2 receptor

Previously, we have shown that sigma-2R/TMEM97 regulates the internalization of lipoproteins via its interaction with the LDLR, acting as a protein that enhances the receptor’s recognition and subsequent endocytosis of its ligands ^6,7^. This raised the possibility that sigma-2R/TMEM97 may regulate SARS-CoV-2 uptake via its interaction with its receptor ACE2, either by enhancing the capacity for this receptor to recognize SARS-CoV-2 directly, or by enhancing endocytosis of the virus upon recognition of the receptor. To assess this possibility, we performed a proximity ligation assay (PLA) on scramble/cas9 cells that were transduced with ACE2. PLA data showed that TMEM97 and ACE2 do indeed interact (Fig 4), forming a protein complex. These findings provided a rationale for the role that sigma-2R/TMEM97 plays in SARS-CoV-2 viral entry into the cell through the ACE2 receptor. In the presence of sigma-2R/TMEM97, the protein complex is intact and facilitates enhanced viral uptake, however upon disruption of this complex the viral uptake was attenuated. PLA signal was absent in scramble/cas9 cells that were not transduced with ACE2 and with sigma-2R/TMEM97 KO cells that were transduced with ACE2 (SI Appendix Fig S1), indicating that if either protein is absent then no interaction was detected. This indicates that sigma-2R/TMEM97 forms a complex with ACE2 and modulates its ability to internalize the SARS-CoV-2 pseudovirus as we have previously observed with the LDLR.

**Figure 4.**
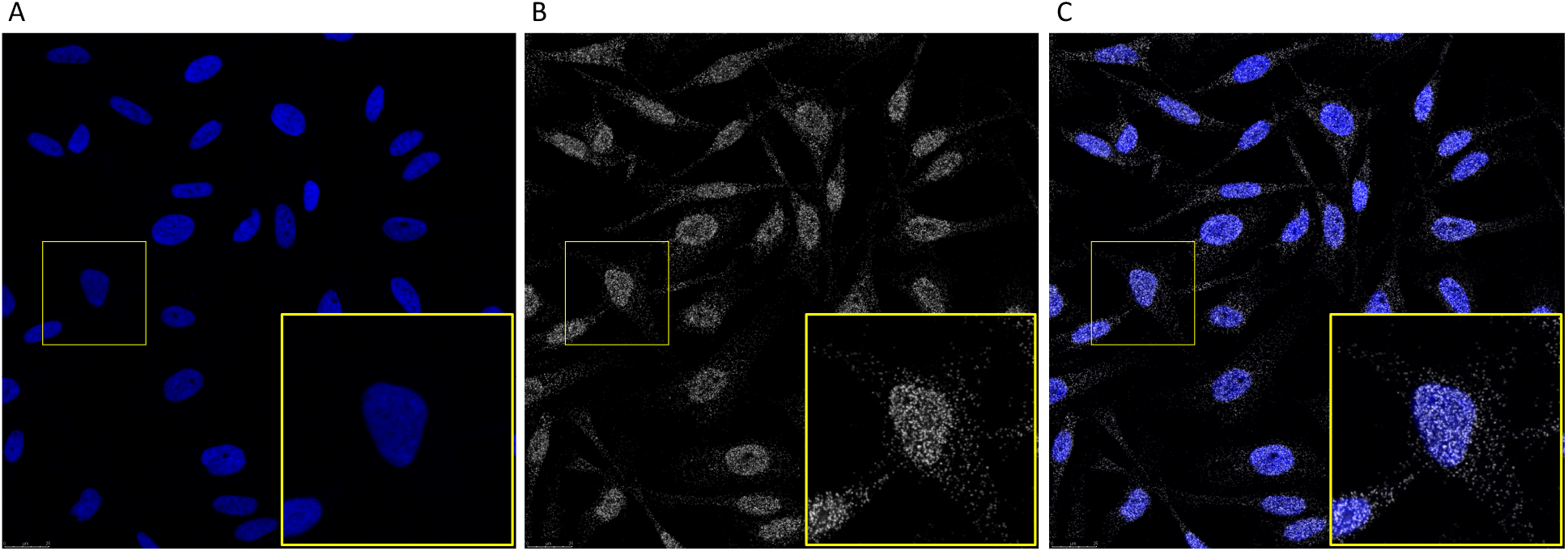
Proximity ligation assay shows TMEM97 interacts with ACE2. HeLa Scramble/Cas9 cells were transduced with human ACE2 expressing an RFP tag. Fluorescent signals for (A) DAPI, (B) PLA signal, (C) merged overlay indicated the presence of a protein complex involving TMEM97 and ACE2. Area of interested indicated has been enlarged to show detail (insert). (n=3)

## DISUCSSION

In this study, we first aimed to identify genes that are differentially expressed in sigma-2R/TMEM97 KO cells. Amongst the top upregulated genes were C4BPB (logFC = 3.33) and C4BPA (logFC = 2.20), which are proteins responsible for regulating the complement cascade upon tissue damage and modulating the coagulation cascade through its interaction with protein S ^24,34^. Previous studies have shown the importance of inhibition of C4BP in the context of pathogenic infections including IAV and ebola virus ^27,37^. Our next step was to determine if the protein expression levels tracked the genetic expression levels of C4BP upregulation upon loss of sigma-2R/TMEM97. Results indicated that cells exhibited a significant increase in secretion of C4BP as determined by ELISA (Fig 2A) upon knocking out sigma-2R/TMEM97 and treatment with sigma-2R/TMEM97 specific ligand RHM-4 in a dose dependent manner (Fig 2B). The observation that C4BP expression was observed at both the transcriptional and translational levels indicated a novel function of sigma-2R/TMEM97 in regulating the complement cascade, indicating a novel function of the protein in inflammation and thrombosis, furthering our understanding of the biological role of sigma-2R/TMEM97 beyond its role in the endocytic pathway.

It is likely that inhibition of sigma-2R/TMEM97 either through knockout or treatment with small molecules affect viral entry into the cell via its interaction with the ACE2 receptor directly, or indirectly through the modulation of the complement pathway or via C4BP upregulation. To delineate this, we used HeLa sigma-2R/TMEM97 KO cells that were previously generated in our lab and treated them with a pseudovirus that expresses the SARS-CoV-2 spike protein. These cells displayed the decreased capacity of SARS-CoV-2 pseudovirus viral entry into the cell, indicating the importance of sigma-2R/TMEM97 in interacting with host proteins to facilitate pathogenic uptake. We also observed that treatment with haloperidol, a sigma receptor ligand resulted in the decreased uptake of pseudovirus. Moreover, RHM-4, which is specific to the sigma-2R/TMEM97 also decreased the pseudovirus uptake, indicating that the inhibition of sigma-2R/TMEM97 specifically is a novel strategy to reduce viral uptake.

In this study, we have shown that sigma-2R/TMEM97 directly interacts with ACE2 receptor, which is analogous to our previous findings that sigma-2R/TMEM97 is involved with facilitating effective uptake of ligands such as LDL, apoE, and AB42 through its interaction with the LDLR directly. It is likely that inhibition of sigma-2R/TMEM97 results in the disruption of the sigma-2R/TMEM97-ACE2 protein complex, and subsequent increases in C4BP levels facilitate an environment that results in inhibition of the endocytic process involving SARS-CoV-2 recognition of the ACE2 receptor and or downstream endocytic machinery allowing the entry of the virus into the cells ^13^. Moreover, this effect on inhibition of the complement cascade through the reduction of C4BP may prove to have a beneficial anti-inflammatory and anti-thrombotic effect on patients infected by the SARS-CoV-2 virus. Taken together this data indicates that the sigma-2R/TMEM97 has been implicated with pathogen infection of host cells, further expanding the known biological entities that sigma-2R/TMEM97 is involved with entry into cells beyond endogenous ligands such as lipoproteins or intrinsically disordered proteins. The pharmacological targeting of sigma-2R/TMEM97 has thus far shown to be beneficial in a variety of diseases including Alzheimer’s Disease, the dysregulation of lipid homeostasis, and now viral infectivity, through its interaction with host receptors and modulation of endocytic pathway. These observations indicate the importance of the receptor in the uptake pathway and hint towards its potential role as a modulating protein enhancing receptor function. Furthermore, we have shown a unique biological role of sigma-2R/TMEM97 in regulating the inflammatory pathway.

The similarity between the studies involving CRISPR/cas9 ablation of sigma-2R/TMEM97 and pharmacological inhibition of the protein further support the idea that sigma-2R/TMEM97 is a potential therapeutic target for decreasing effective viral uptake which would result decreased infectivity, helping to curb the transmission of the virus within the population during a pandemic. While vaccines have been developed against SARS-CoV-2 viruses, having therapeutics that target the biology of host cells may be important to limit the infectivity of new variants of the constantly mutating virus and of future coronoaviruses where current vaccines may not prove to be effective ^38^.

In conclusion, we have shown that sigma-2R/TMEM97 regulates the effectiveness of SARS-CoV-2 pseudovirus entry in a targetable manner by modulating their entry into the cells, potentially due to its interaction with the ACE2 receptor and that inhibition of sigma-2R/TMEM97 may prove to have therapeutic value through downstream effects on the complement and thrombotic cascade. These observations provide an interesting candidate to be developed into a therapeutic that can be used to inhibit viral uptake and subsequent inflammatory effects. Small molecules targeting sigma-2R/TMEM97 are likely to be excellent candidates for prophylactic therapeutic development against SARS-CoV-2 uptake and subsequent infection, and future work will reveal the importance of sigma-2R/TMEM97 in other types of pathogenic infections.

## Methods

### Materials

*N*-(4-(6,7-dimethoxy-3,4-dihydroisoquinolin-2(1H)-yl)butyl)-2,3-dimethoxy-5-iodo-benzamide and [125I]*N*-(4-(6,7-dimethoxy-3,4-dihydroisoquinolin-2(1H)-yl)butyl)-2,3-dimethoxy-5-iodo-benzamide were synthesized as previously described ^39,40^. Hoechst 33342 (BD Pharmingen, 561908).

### Cell Culture

HeLa cell sigma-2R/TMEM97 knockout cell lines were generated as previously described ^6,7^. Cells were cultured in MEM with 10% FBS, 1X penicillin/streptomycin, 2mM L-glutamine, and 1X MEM non-essential amino acids. For uptake experiments, cells were plated, and incubated for 24 hours, media was removed and cells were incubated in MEM containing 10% lipoprotein depleted serum for an additional 24 hours prior to treatment.

### RNA-Seq

Total RNA was extracted from sigma2-R/TMEM97 KO Hela cells or scramble/cas9 control cells in triplicates. RNAseq libraries were generated. Libraries were sequenced by University of Pennsylvania Next-Generation Sequencing Core on an Illumina HiSeq4000 sequencer (sequencer mode: Hight Output4K; serial number: Vn:K00315; Run Time Analysis Software version: 2.7.7; HiSeq control software version: HD 3.4.0.38; Run mode: 150PE). FASTQ files were aligned to the human genome (hg19) using STAR aligner ^41^ (version 2.7.1a) requiring the marking of multi-mappers and duplicates unique mappers (--bamRemoveDuplicatesType UniqueIdentical). Impact of sigma2-R/TMEM97 KO on HeLa cells transcriptome was assessed using Rsubread / edgeR pipeline (version 2.4.2 and 3.32.1, respectively) ^42^. Paired-end fragments were counted at the meta-feature level (*i*.*e*., genes) with the featuresCounts utility ^43^ implemented in Rsubread, using in-built human genome reference annotation (NCBI RefSeq hg19), requiring duplicate reads to be ignored and successful alignment of both ends from the same fragment before assignment of the fragment to a meta-feature. Read counts matrix was filtered to remove genes with less than 1 read per kilobase per million mapped fragments (RPKM) and library sizes were normalized using edgeR calcNomFactor function which implements a trimmed mean of M values normalization method ^44^. Differential gene expression between TMEM97 KO cells and control cells was assessed using edgeR gene-wise negative binomial generalized linear model with quasi-likelihood testing method (glmQLfit).

### C4BPB ELISA

HeLa cells were plated in 6 well plates at 100,000 cells/well for 72 hours in normal media in the presence or absence of drug treatment as indicated. 72 hours later, media was collected and C4BPB was quantified via ELISA according to manufacturer’s instructions (Abcam, ab222866).

### ACE2 Transduction and SARS-CoV-2 Pseudovirus

HeLa cells were plated in 96 well plates at 10,000 cells/well, and transduced with human ACE2 with an RFP tag according to manufacturer (Montana Molecular #C1100R). 24 hours later cells were incubated with pseudovirus expressing SARS-CoV-2 spike protein according to manufacturer (Montan Molecular #C1110G). Fluorescent signal was detected using a Perkin Elmer Enspire® Multimode Plate Reader. For drug treatment, signal was normalized to cells that were not treated with compounds. IC50 was determined using GraphPad Prism 9.0.

### Confocal Microscopy and Proximity Ligation Assay

Cells were plated in 8-well chamber slides (Lab-Tek cc2 plates, 154534PK Thermo Scientific) and transduced with human ACE2 with an RFP tag according to manufacturer (Montana Molecular #C1100R). Cells were washed 3 times with PBS then fixed with 4% paraformaldehyde (Santa Cruz) for 10 minutes at room temperature, washed 3 times with PBS, then permeabilized with 0.1% Triton X-100 in PBS for 10 minutes at room temperature. Cells were blocked with 10% Goat Serum (50062Z Thermo Scientific) for one hour then incubated with rabbit anti-TMEM97 primary antibody (Novus NBP1-30436) 1:200 in PLA buffer diluent and mouse anti-RFP (Invitrogen MA515257) overnight, washed 3 times with PLA Wash buffer B 3 times, proceeded with PLA assay according to manufacturer (Duolink In Situ PLA Far Red kit Sigma DUO92105). Cells were then mounted and images were acquired at 40X magnification on a Leica STED 8X Super-resolution Confocal Microscope.

## Notes

### Competing Interest Statement

The authors have declared no competing interest.

## References

1. Martin, W. R., Eades, C. G., Thompson, J. A., Huppler, R. E., Gilbert, P. E., The effects of morphine- and nalorphine-like drugs in the nondependent and morphine-dependent chronic spinal dog. The Journal of pharmacology and experimental therapeutics 1976, 197 (3), 517–32.

2. Hellewell, S. B., Bruce, A., Feinstein, G., Orringer, J., Williams, W., Bowen, W. D., Rat liver and kidney contain high densities of sigma 1 and sigma 2 receptors: characterization by ligand binding and photoaffinity labeling. European journal of pharmacology 1994, 268 (1), 9–18.

3. Matsumoto, R. R., σ Receptors: Historical Perspective and Background. In Sigma Receptors: Chemistry, Cell Biology and Clinical Implications, Su, T.-P., Matsumoto, R. R., Bowen, W. D., Eds. Springer US: Boston, MA, 2007; pp 1–23.

4. Xu, J., Zeng, C., Chu, W., Pan, F., Rothfuss, J. M., Zhang, F., Tu, Z., Zhou, D., Zeng, D., Vangveravong, S., Johnston, F., Spitzer, D., Chang, K. C., Hotchkiss, R. S., Hawkins, W. G., Wheeler, K. T., Mach, R. H., Identification of the PGRMC1 protein complex as the putative sigma-2 receptor binding site. Nature Communications 2011, 2, 380.

5. Alon, A., Schmidt, H. R., Wood, M. D., Sahn, J. J., Martin, S. F., Kruse, A. C., Identification of the gene that codes for the sigma2 receptor. Proceedings of the National Academy of Sciences of the United States of America 2017, 114 (27), 7160–7165.

6. Riad, A., Zeng, C., Weng, C. C., Winters, H., Xu, K., Makvandi, M., Metz, T., Carlin, S., Mach, R. H., Sigma-2 Receptor/TMEM97 and PGRMC-1 Increase the Rate of Internalization of LDL by LDL Receptor through the Formation of a Ternary Complex. Sci Rep 2018, 8 (1), 16845.

7. Riad, A., Lengyel-Zhand, Z., Zeng, C., Weng, C. C., Lee, V. M., Trojanowski, J. Q., Mach, R. H., The Sigma-2 Receptor/TMEM97, PGRMC1, and LDL Receptor Complex Are Responsible for the Cellular Uptake of Abeta42 and Its Protein Aggregates. Mol Neurobiol 2020, 57 (9), 3803–3813.

8. Ebrahimi-Fakhari, D., Wahlster, L., Bartz, F., Werenbeck-Ueding, J., Praggastis, M., Zhang, J., Joggerst-Thomalla, B., Theiss, S., Grimm, D., Ory, D. S., Runz, H., Reduction of TMEM97 increases NPC1 protein levels and restores cholesterol trafficking in Niemann-pick type C1 disease cells. Hum Mol Genet 2016, 25 (16), 3588–3599.

9. Bartz, F., Kern, L., Erz, D., Zhu, M., Gilbert, D., Meinhof, T., Wirkner, U., Erfle, H., Muckenthaler, M., Pepperkok, R., Runz, H., Identification of cholesterol-regulating genes by targeted RNAi screening. Cell metabolism 2009, 10 (1), 63–75.

10. Gordon, D. E., Jang, G. M., Bouhaddou, M., Xu, J., Obernier, K., White, K. M., O’Meara, M. J., Rezelj, V. V., Guo, J. Z., Swaney, D. L., Tummino, T. A., Huttenhain, R., Kaake, R. M., Richards, A. L., Tutuncuoglu, B., Foussard, H., Batra, J., Haas, K., Modak, M., Kim, M., Haas, P., Polacco, B. J., Braberg, H., Fabius, J. M., Eckhardt, M., Soucheray, M., Bennett, M. J., Cakir, M., McGregor, M. J., Li, Q., Meyer, B., Roesch, F., Vallet, T., Mac Kain, A., Miorin, L., Moreno, E., Naing, Z. Z. C., Zhou, Y., Peng, S., Shi, Y., Zhang, Z., Shen, W., Kirby, I. T., Melnyk, J. E., Chorba, J. S., Lou, K., Dai, S. A., Barrio-Hernandez, I., Memon, D., Hernandez-Armenta, C., Lyu, J., Mathy, C. J. P., Perica, T., Pilla, K. B., Ganesan, S. J., Saltzberg, D. J., Rakesh, R., Liu, X., Rosenthal, S. B., Calviello, L., Venkataramanan, S., Liboy-Lugo, J., Lin, Y., Huang, X. P., Liu, Y., Wankowicz, S. A., Bohn, M., Safari, M., Ugur, F. S., Koh, C., Savar, N. S., Tran, Q. D., Shengjuler, D., Fletcher, S. J., O’Neal, M. C., Cai, Y., Chang, J. C. J., Broadhurst, D. J., Klippsten, S., Sharp, P. P., Wenzell, N. A., Kuzuoglu-Ozturk, D., Wang, H. Y., Trenker, R., Young, J. M., Cavero, D. A., Hiatt, J., Roth, T. L., Rathore, U., Subramanian, A., Noack, J., Hubert, M., Stroud, R. M., Frankel, A. D., Rosenberg, O. S., Verba, K. A., Agard, D. A., Ott, M., Emerman, M., Jura, N., von Zastrow, M., Verdin, E., Ashworth, A., Schwartz, O., d’Enfert, C., Mukherjee, S., Jacobson, M., Malik, H. S., Fujimori, D. G., Ideker, T., Craik, C. S., Floor, S. N., Fraser, J. S., Gross, J. D., Sali, A., Roth, B. L., Ruggero, D., Taunton, J., Kortemme, T., Beltrao, P., Vignuzzi, M., Garcia-Sastre, A., Shokat, K. M., Shoichet, B. K., Krogan, N. J., A SARS-CoV-2 protein interaction map reveals targets for drug repurposing. Nature 2020, 583 (7816), 459–468.

11. Hoffmann, M., Kleine-Weber, H., Schroeder, S., Kruger, N., Herrler, T., Erichsen, S., Schiergens, T. S., Herrler, G., Wu, N. H., Nitsche, A., Muller, M. A., Drosten, C., Pohlmann, S., SARS-CoV-2 Cell Entry Depends on ACE2 and TMPRSS2 and Is Blocked by a Clinically Proven Protease Inhibitor. Cell 2020, 181 (2), 271–280 e8.

12. Dominguez Andres, A., Feng, Y., Campos, A. R., Yin, J., Yang, C.-C., James, B., Murad, R., Kim, H., Deshpande, A. J., Gordon, D. E., Krogan, N., Pippa, R., Ronai, Z. e. A., SARS-CoV-2 ORF9c Is a Membrane-Associated Protein that Suppresses Antiviral Responses in Cells. bioRxiv 2020, 2020.08.18.256776.

13. Fletcher-Sandersjoo, A., Bellander, B. M., Is COVID-19 associated thrombosis caused by overactivation of the complement cascade? A literature review. Thromb Res 2020, 194, 36–41.

14. Wiersinga, W. J., Rhodes, A., Cheng, A. C., Peacock, S. J., Prescott, H. C., Pathophysiology, Transmission, Diagnosis, and Treatment of Coronavirus Disease 2019 (COVID-19): A Review. JAMA 2020, 324 (8), 782–793.

15. Yu, J., Yuan, X., Chen, H., Chaturvedi, S., Braunstein, E. M., Brodsky, R. A., Direct activation of the alternative complement pathway by SARS-CoV-2 spike proteins is blocked by factor D inhibition. Blood 2020, 136 (18), 2080–2089.

16. Leonard, B. E., Sigma receptors and sigma ligands: background to a pharmacological enigma. Pharmacopsychiatry 2004, 37 Suppl 3, S166–70.

17. Izzo, N. J., Staniszewski, A., To, L., Fa, M., Teich, A. F., Saeed, F., Wostein, H., Walko, T., 3rd; Vaswani, A., Wardius, M., Syed, Z., Ravenscroft, J., Mozzoni, K., Silky, C., Rehak, C., Yurko, R., Finn, P., Look, G., Rishton, G., Safferstein, H., Miller, M., Johanson, C., Stopa, E., Windisch, M., Hutter-Paier, B., Shamloo, M., Arancio, O., LeVine, H., 3rd; Catalano, S.M., Alzheimer’s therapeutics targeting amyloid beta 1-42 oligomers I: Abeta 42 oligomer binding to specific neuronal receptors is displaced by drug candidates that improve cognitive deficits. PloS one 2014, 9 (11), e111898.

18. Izzo, N. J., Xu, J., Zeng, C., Kirk, M. J., Mozzoni, K., Silky, C., Rehak, C., Yurko, R., Look, G., Rishton, G., Safferstein, H., Cruchaga, C., Goate, A., Cahill, M. A., Arancio, O., Mach, R. H., Craven, R., Head, E., LeVine, H., 3rd; Spires-Jones, T. L., Catalano, S. M., Alzheimer’s therapeutics targeting amyloid beta 1-42 oligomers II: Sigma-2/PGRMC1 receptors mediate Abeta 42 oligomer binding and synaptotoxicity. PloS one 2014, 9 (11), e111899.

19. Mach, R. H., Smith, C. R., al-Nabulsi, I., Whirrett, B. R., Childers, S. R., Wheeler, K. T., Sigma 2 receptors as potential biomarkers of proliferation in breast cancer. Cancer Res 1997, 57 (1), 156–61.

20. Mach, R. H., Zeng, C., Hawkins, W. G., The sigma2 receptor: a novel protein for the imaging and treatment of cancer. Journal of medicinal chemistry 2013, 56 (18), 7137–60.

21. McDonald, E. S., Doot, R. K., Young, A. J., Schubert, E. K., Tchou, J., Pryma, D. A., Farwell, M. D., Nayak, A., Ziober, A., Feldman, M. D., DeMichele, A., Clark, A. S., Shah, P. D., Lee, H., Carlin, S. D., Mach, R. H., Mankoff, D. A., Breast Cancer (18)F-ISO-1 Uptake as a Marker of Proliferation Status. J Nucl Med 2020, 61 (5), 665–670.

22. Wheeler, K. T., Wang, L. M., Wallen, C. A., Childers, S. R., Cline, J. M., Keng, P. C., Mach, R. H., Sigma-2 receptors as a biomarker of proliferation in solid tumours. Br J Cancer 2000, 82 (6), 1223–32.

23. Zeng, C., Riad, A., Mach, R. H., The Biological Function of Sigma-2 Receptor/TMEM97 and Its Utility in PET Imaging Studies in Cancer. Cancers (Basel) 2020, 12 (7).

24. Ermert, D., Blom, A. M., C4b-binding protein: The good, the bad and the deadly. Novel functions of an old friend. Immunol Lett 2016, 169, 82–92.

25. Fletcher-Sandersjoo, A., Maegele, M., Bellander, B. M., Does Complement-Mediated Hemostatic Disturbance Occur in Traumatic Brain Injury? A Literature Review and Observational Study Protocol. Int J Mol Sci 2020, 21 (5).

26. Mohlin, F. C., Blom, A. M., Purification and functional characterization of C4b-binding protein (C4BP). Methods Mol Biol 2014, 1100, 169–76.

27. Varghese, P. M., Murugaiah, V., Beirag, N., Temperton, N., Khan, H. A., Alrokayan, S. H., Al-Ahdal, M. N., Nal, B., Al-Mohanna, F. A., Sim, R. B., Kishore, U., C4b Binding Protein Acts as an Innate Immune Effector Against Influenza A Virus. Front Immunol 2020, 11, 585361.

28. Bilaloglu, S., Aphinyanaphongs, Y., Jones, S., Iturrate, E., Hochman, J., Berger, J. S., Thrombosis in Hospitalized Patients With COVID-19 in a New York City Health System. JAMA 2020, 324 (8), 799–801.

29. Connors, J. M., Levy, J. H., COVID-19 and its implications for thrombosis and anticoagulation. Blood 2020, 135 (23), 2033–2040.

30. Java, A., Apicelli, A. J., Liszewski, M. K., Coler-Reilly, A., Atkinson, J. P., Kim, A. H., Kulkarni, H. S., The complement system in COVID-19: friend and foe? JCI Insight 2020, 5 (15).

31. Klok, F. A., Kruip, M., van der Meer, N. J. M., Arbous, M. S., Gommers, D., Kant, K. M., Kaptein, F. H. J., van Paassen, J., Stals, M. A. M., Huisman, M. V., Endeman, H., Incidence of thrombotic complications in critically ill ICU patients with COVID-19. Thromb Res 2020, 191, 145–147.

32. Kulkarni, H. S., Atkinson, J. P., Targeting complement activation in COVID-19. Blood 2020, 136 (18), 2000–2001.

33. Lee, M. H., Perl, D. P., Nair, G., Li, W., Maric, D., Murray, H., Dodd, S. J., Koretsky, A. P., Watts, J. A., Cheung, V., Masliah, E., Horkayne-Szakaly, I., Jones, R., Stram, M. N., Moncur, J., Hefti, M., Folkerth, R. D., Nath, A., Microvascular Injury in the Brains of Patients with Covid-19. N Engl J Med 2020.

34. Criado Garcia, O., Sanchez-Corral, P., Rodriguez de Cordoba, S., Isoforms of human C4b-binding protein. II. Differential modulation of the C4BPA and C4BPB genes by acute phase cytokines. J Immunol 1995, 155 (8), 4037–43.

35. Moffat, G. J., Tack, B. F., Regulation of C4b-binding protein gene expression by the acute-phase mediators tumor necrosis factor-alpha, interleukin-6, and interleukin-1. Biochemistry 1992, 31 (49), 12376–84.

36. Phillips, D. J., Novinger, M. S., Evatt, B. L., Hooper, W. C., TNF-alpha suppresses IL-1 alpha and IL-6 upregulation of C4b-binding protein in HepG-2 hepatoma cells. Thromb Res 1996, 81 (3), 307–14.

37. Ward, M. D., Brueggemann, E. E., Kenny, T., Reitstetter, R. E., Mahone, C. R., Trevino, S., Wetzel, K., Donnelly, G. C., Retterer, C., Norgren, R. B., Jr., Panchal, R. G., Warren, T. K., Bavari, S., Cazares, L. H., Characterization of the plasma proteome of nonhuman primates during Ebola virus disease or melioidosis: a host response comparison. Clin Proteomics 2019, 16, 7.

38. Sharma, O., Sultan, A. A., Ding, H., Triggle, C. R., A Review of the Progress and Challenges of Developing a Vaccine for COVID-19. Front Immunol 2020, 11, 585354.

39. Hou, C., Tu, Z., Mach, R., Kung, H. F., Kung, M. P., Characterization of a novel iodinated sigma-2 receptor ligand as a cell proliferation marker. Nuclear Medicine and Biology 2006, 33 (2), 203–9.

40. Xu, J., Tu, Z., Jones, L. A., Vangveravong, S., Wheeler, K. T., Mach, R. H., [3H]N-[4-(3,4-dihydro-6,7-dimethoxyisoquinolin-2(1H)-yl)butyl]-2-methoxy-5-methyl benzamide: a novel sigma-2 receptor probe. European journal of pharmacology 2005, 525 (1-3), 8–17.

41. Dobin, A., Davis, C. A., Schlesinger, F., Drenkow, J., Zaleski, C., Jha, S., Batut, P., Chaisson, M., Gingeras, T. R., STAR: ultrafast universal RNA-seq aligner. Bioinformatics 2013, 29 (1), 15–21.

42. Chen, Y., Lun, A. T., Smyth, G. K., From reads to genes to pathways: differential expression analysis of RNA-Seq experiments using Rsubread and the edgeR quasi-likelihood pipeline. F1000Res 2016, 5, 1438.

43. Liao, Y., Smyth, G. K., Shi, W., featureCounts: an efficient general purpose program for assigning sequence reads to genomic features. Bioinformatics 2014, 30 (7), 923–30.

44. Robinson, M. D., Oshlack, A., A scaling normalization method for differential expression analysis of RNA-seq data. Genome Biol 2010, 11 (3), R25.

