## Supplementary Figure for "Differential gene expression by RNA-Seq in Sigma-2 Receptor/TMEM97 knockout cells reveals its role in complement activation and SARS-CoV-2 viral uptake"

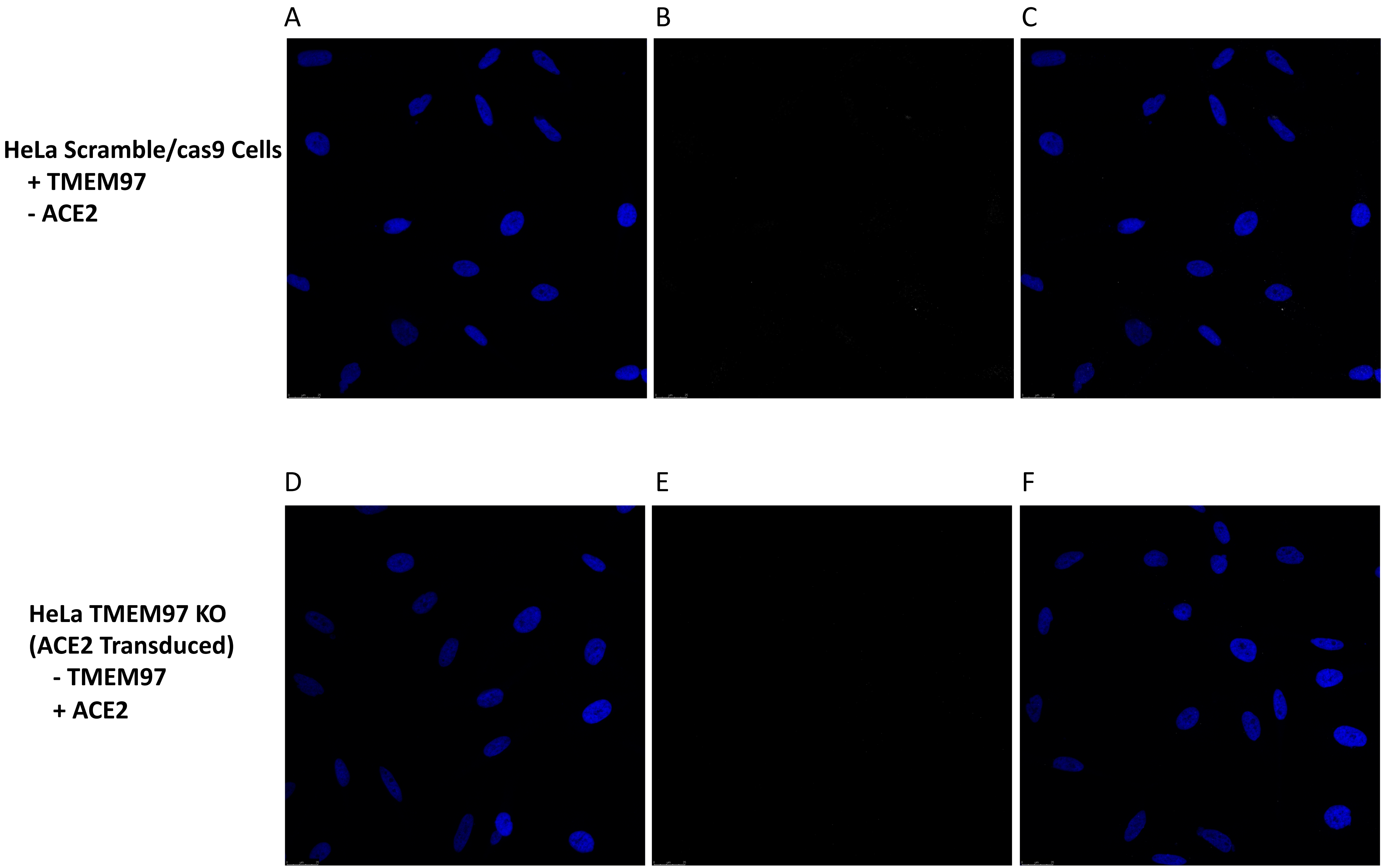


**Supplementary Figure 1.** Proximity ligation assay shows no signals in the absence of TMEM97 or ACE2. Scramble/cas9 cells lacking ACE2 (A-C) and TMEM97 KO cells transduced with ACE2 (D-F) show no interaction in the absence of either protein. (A, D) DAPI, (B, E) PLA signal, (C, F) merged overlay. (n=3)
